# A Bigenic mouse model of FSGS reveals perturbed pathways in podocytes, mesangial cells and endothelial cells

**DOI:** 10.1101/613679

**Authors:** Andrew S. Potter, Keri Drake, Eric W. Brunskill, S. Steven Potter

## Abstract

Focal segmental glomerulosclerosis is a major cause of end stage renal disease. Many patients prove unresponsive to available therapies. An improved understanding of the molecular basis of the disease process could provide insights leading to novel therapeutic approaches. In this study we carried out an RNA-seq analysis of the altered gene expression patterns of podocytes, mesangial cells and glomerular endothelial cells of the bigenic *Cd2ap*+/-, *Fyn*-/- mutant mouse model of FSGS. In the podocytes we observed upregulation of many genes related to the Tgfβ family/pathway, including *Gdnf, Tgfβ1, Tgfβ2, Snai2, Vegfb, Bmp4*, and *Tnc*. The mutant podocytes also showed upregulation of *Acta2*, a marker of smooth muscle and associated with myofibroblasts, which are implicated in driving fibrosis. GO analysis of the podocyte upregulated genes identified elevated protein kinase activity, increased expression of growth factors, and negative regulation of cell adhesion, perhaps related to the observed podocyte loss. Both podocytes and mesangial cells showed strong upregulation of aldehyde dehydrogenase genes involved in the synthesis of retinoic acid. Similarly, the *Cd2ap*+/-, *Fyn*-/- mesangial cells, as well as podocytes in other genetic models, and the glomeruli of human FSGS patients, all show upregulation of the serine protease Prss23, with the common thread suggesting important functionality. Another gene with strong upregulation in the *Cd2ap*+/-, *Fyn*-/- mutant mesangial cells as well as multiple other mutant mouse models of FSGS was thrombospondin, which activates the secreted inactive form of Tgfβ. The *Cd2ap*+/-, *Fyn*-/- mutant endothelial cells showed elevated expression of genes involved in cell proliferation, angioblast migration, angiogenesis, and neovasculature, all consistent with the formation of new blood vessels in the diseased glomerulus. The resulting global definition of the perturbed molecular pathways in the three major cell types of the mutant glomerulus provide deeper understanding of the molecular pathogenic pathways.

## Introduction

Focal segmental glomerulosclerosis (FSGS) is a histologic pattern that is the most common glomerular cause of end stage renal disease (ESRD) in the United States (1). It is characterized by sclerosis of parts (segmental) of some (focal) glomeruli. There is typically evidence of collapse of capillary loops and increased mesangial matrix deposition in segments of a portion of glomeruli. FSGS histologic lesion is found in about 35% of all cases, and 50% of African Americans, with nephrotic syndrome (2). There is a trend of increasing incidence of FSGS caused ESRD in the United States (3) and world (4).

The Columbia morphological classification defines the collapsing variant, tip variant, perihilar variant, cellular variant and classical or NOS (not otherwise specified) FSGS (5, 6). It is also possible to cluster FSGS into six clinical types based on etiology (7), which includes the less common medication or infection associated FSGS. Adaptive FSGS results from glomerular hyperfiltration, which can be associated with low birth weight, morbid obesity or sickle cell anemia. Another type is idiopathic, or primary FSGS, with no known cause. In addition, one can define two types of FSGS with a clear genetic basis. First, there are high penetrance genetic causes, with over 38 genes now identified, where homozygous mutation of a single gene results in FSGS (7). These are more likely seen in childhood nephrotic syndrome (∼60%) than in older children or adolescents (∼5%), with still lower rates in adults (8, 9). Causative genes include *Actn4, Inf2, Trpc6*, and *Nphs2*, to name a few (7).

A second genetic cause category involves multiple mutant allele combinations that contribute to FSGS with lower penetrance. For example, genetic variants of *Apol1* are a major contributing factor to FSGS in individuals of sub-Saharan descent, being associated with 72% of cases (10). The effect is mostly recessive, with two risk alleles required, but penetrance is low, as most individuals with two risk alleles will not develop FSGS. Presumably additional environmental and/or genetic contributions are required. Indeed, it is generally thought that monogenic disease is relatively rare compared to multifactorial (multiple mutant genes combined with environmental causes) and polygenic (mutations in multiple genes) disease. The cumulative effects of several mutations in different genes can combine to cause FSGS or modulate its severity. For example, homozygous MYO1E mutation is associated with childhood FSGS (11), while coinheritance of mutations in both COl4A5 and MYO1E can dramatically accentuate disease severity (12).

It has also been shown in mouse models that there can be combined polygenic contributions to FSGS. Cd2ap is a scaffold protein located in the slit diaphragms of podoctyes where it interacts with nephrin and podocin (13, 14). Homozygous mutation of *Cd2ap* has been shown to cause high penetrance FSGS in humans (15, 16). Mice with homozygous mutation of *Cd2ap* also develop FSGS like disease, with severe nephrotic syndrome, extracellular matrix deposition, glomerulosclerosis, extensive podocyte foot process effacement, and death within weeks of birth (13). The phenotype of heterozygous mice with only one *Cd2ap* mutation, however, is “relatively unremarkable” (17), with some glomerular changes noted at 9 months of age (18). *Fyn* encodes a tyrosine kinase, related to *Src*, that phosphorylates the slit diaphragm component Nephrin (19). Heterozygous mutation of *Fyn* gives rise to very rare proteinuria, while homozygous *Fyn* mutation results in proteinuria in only 31% of mice at an average onset of 8 months (17). Of interest, however, combined *Cd2ap*+/-, *Fyn*-/- mice develop proteinuria in 100% of mice at average onset of only 5 months (17). This synergy of phenotype represents a combined susceptibility allele model. A fraction of human FSGS cases also likely result from multiple mutant alleles, in various mixes of susceptibility genes, together increasing the chances of FSGS. In this report we further study the *Cd2ap*+/-, *Fyn*-/- bigenic murine model of FSGS, examining gene expression changes that take place in the glomerular podocyte, mesangial and endothelial cells.

Podocyte injury plays a key role in the initiation and progression of glomerular disease, including FSGS (20). The first detectable morphological characteristics of FSGS are found in podocytes (21), which undergo hypertrophy, detachment from the glomerular basement membrane (GBM), effacement of foot processes and depletion in number. The other two main glomerular cell types, however, the mesangial and endothelial cells, also undergo dramatic changes during FSGS. There is, for example, mesangial expansion with increased extracellular matrix, and endothelial cells can show increased leukocyte recruitment (22) as well as *de novo* angiogenesis, which can result in leaky vessels (23). A comprehensive analysis of FSGS, therefore, requires examination of mesangial cells and endothelial cells as well as podocytes.

The current Kidney Disease: Improving Global Outcome (KDIGO) practice guidelines link therapy to pathology. Initial treatments include inhibitors of the renin-angiotensin system and corticosteroids. Steroid resistant patients can be treated with cyclosporine, mycophenolate mofetil, or tacrolimus, with responses varying for different types of FSGS. Nevertheless, a high percentage of patients prove unresponsive to all available therapies, emphasizing the need for a deeper understanding of FSGS to guide the development of improved treatment options. In this report we define the activated pathogenic and protective molecular pathways in each major cell type of the glomerulus in the bigenic *Cd2ap*+/-, *Fyn*-/- mouse model of FSGS, thereby providing a global view the disease process that might aid in the identification of novel therapeutic targets.

## Materials and Methods

### Mouse Strains

The *Cd2ap* mutant (B6.129X1-*Cd2ap*^tm1Shaw^/J), *Tie2-GFP* (Tg[*TIE2GFP*]287Sato/J) and Fyn (B6.129-*Fyn*^tm1Sor^/J) mice were obtained from the Jackson Laboratory. *MafB-GFP*, Tg (*MafB-EGFP)* FT79Gsat and *Meis1-GFP* Tg (*Meis1-EGFP*) FO156Gsat, were from GENSAT/MMRC (http://www.gensat.org).

### Animal Ethics

All animal experiments were carried out according to protocols approved by the Cincinnati Children’s Medical Center Institutional Animal Care and Use Committee (protocol title “mouse models of focal segmental glomerulosclerosis”, number 2016-0020).

### Single Cell Prep for FACS Sorting

Mice were euthanized via cervical dislocation after sedation with isoflurane; kidneys were isolated and placed into ice-cold PBS. Glomeruli were isolated as previously described (24).

For isolating mesangial and endothelial cells, the glomeruli were pelleted, rinsed with ice-cold PBS, and then re-suspended in 200 µL TrypLE Select 10x, incubated at 37° C for 10-15 minutes, and triturated vigorously every 3 minutes. Cell digestion was monitored by taking a small aliquot and visualizing with a microscope. After digestion ice-cold 10% FBS/PBS was added and the cells were triturated vigorously. The cells were then pelleted, washed with ice-cold 1% FBS/PBS and filtered using a 35-µM filter mesh (Falcon, cat. # 352235) before FACS sorting. GFP positive cells were collected into lysis buffer containing 0.1% SDS (in H_2_0). After collection, the lysate was vortexed and placed on dry ice.

For isolating podocytes, a single cell suspension was derived from the glomeruli as previously described (24). Briefly, the glomeruli were incubated for 40 minutes in 2 mL enzymatic digest buffer (Containing Type 2 collagenase 300 U/mL, 1 mg/mL pronase E, 50 U/mL DNAse 1) at 37° C while shaking at 1400 RPM/min in a thermomixer. Every 10 minutes the digestion mix was passaged two times with a 27-gauge needle (25). After digestion, equal volume ice-cold 10% FBS/PBS was added and the cells were triturated. The cells were filtered using a 40 µM filter to remove clumps, pelleted by centrifugation, rinsed with ice-cold 1% FBS/PBS, re-suspended in 400 µL 1% FBS/PBS and filtered using a 35-µM filter mesh prior to FACS sorting. GFP-positive cells were collected in Buffer RL. After collection, the lysate was vortexed and placed on dry ice.

### Target Cell Isolation

The *Meis1-GFP, MafB-GFP*, and *Tie2-GFP* transgene reporters enabled FACS-sorting purification of mesangial cells, podocytes and endothelial cells, respectively, from single-cell suspensions derived from the glomeruli of control (wild type or one-allele mice), and *Cd2ap*^*+/-*^ *Fyn*^*-/-*^ (3-allele) mice. Although 3-allele *Cd2ap*^*+/-*^ *Fyn*^*-/-*^ mice developed albuminuria at 5 months of age, both the 3-allele and control mice were sacrificed at an average age of approximately 10-14 months, which coincided with 3-allele mice having significantly elevated BUN and increased pathological evidence of FSGS compared to control mice. From 5-9 months of age, the average BUN of 3-allele mice was 29.13 ± 1.2 compared to 26.46 ± 0.97 for control mice. From 10-14 months of age, the average BUN of 3-allele mice was 35.98 ± 2.9 compared to 27.22 ± 1.4 for control mice. The mice sacrificed were all adult (>= 5 months). The first two mice, aged 5 months, (Mesangial cells: 3-allele and control) that we sacrificed did not show substantial differences in the RNA-Seq gene profiles, so subsequently we used older mice ranging in age from 8 months to 1.5 years that showed significant proteinuria as measured by a protein gel. The average age for 3-allele and control mice was as follows in Table 1.

**Table 1.** Average ages of mice used for analysis.

| Cell Type | Control (Age, months) | 3-allele (Age, months) |
| --- | --- | --- |
| Podocyte | 12 | 12 |
| Mesangial | 10 | 11 |
| Endothelial | 14 | 13 |

### RNA Purification, Amplification and Sequencing

For mesangial and endothelial samples, we purified the RNA using Zymo ZR MicroPrep Kit (cat. # R1061) with in-solution DNAse treatment. Due to a lower yield of cells for 3-allele MafB-GFP selected podocytes (due to podocyte cell loss), we used the Norgen Single Cell RNA Purification Kit (cat. # 51800) with on-column DNAse treatment for both the podocyte control and 3-allele samples.

The initial amplification step for all samples was done with the NuGEN Ovation RNA-Seq System v2. The assay was used to amplify RNA samples to create double stranded cDNA. The concentrations were measured using the Qubit dsDNA BR assay. Libraries were then created for all samples. Specifically, the Illumina protocol, the Nextera XT DNA Sample Preparation Kit, was used to create DNA library templates from the double stranded cDNA. The concentrations were measured using the Qubit dsDNA HS assay. The size of the libraries for each sample was measured using the Agilent HS DNA chip. The samples were placed in a pool. The concentration of the pool is optimized to acquire at least 15-20 million reads per sample. Sequencing was performed on the Illumina HiSeq2500, single-end 75 base pair. Sequencing data is available in the Gene Expression Omnibus (GEO), accession number GSE123179.

### RT-qPCR Validations

Glomeruli were isolated from 3-allele and control mice using the sieving method described previously (24). Glomeruli were then lysed by addition of Buffer RLT, vortexing and then placing the tubes on dry ice. RNA was purified using the Qiagen RNEasy Micro Kit (cat. # 74004). RNA was quantified using a NanoDrop spectrophotometer (NanoDrop Technologies, DE, USA). The quality of the RNA was analyzed using an Agilent Bioanalyzer (nano-chip). RNA Integrity Numbers (RIN) were > 8.0. Reverse transcription reactions were set up using the RT-VILO SuperScript cDNA synthesis kit from Invitrogen (cat. # 11754-050). The cDNA was then precipitated by adding 2 µL of 3 M sodium acetate, 1 µL glycogen and 57.5 µL 100% ETOH for a total volume of 80.5 µL. Following overnight incubation at −20 °C, the cDNA was pelleted at 14,000 RPM for 20 minutes at 4° C. The pellet was rinsed with 70% ETOH, decanted and left to air dry. The cDNA was re-suspended in 51 µL H_2_0 and then quantified using a NanoDrop spectrophotometer set to measure single-stranded DNA. RT-qPCR was performed using the PowerUp SYBR Green Master Mix (Applied Biosystems, cat. # A25742) and the StepOnePlus using the relative standard curve method. Samples were normalized to beta-actin (mesangial and endothelial genes) or nephrin (podocyte genes).

### RT-qPCR Primer Design

Genes were selected based on restricted up-regulation in one sorted glomerular 3-allele cell type compared to control cells, or in the case of Serpine1 up-regulation in several cell-types. RT-qPCR primers were designed using Primer-Blast to amplify a product size between 75 and 200 bp using the mRNA RefSeq database of *Mus musculus*. Where possible, the primers were designed to span an intron. Primers used are listed below in Table 2.

**Table 2.**
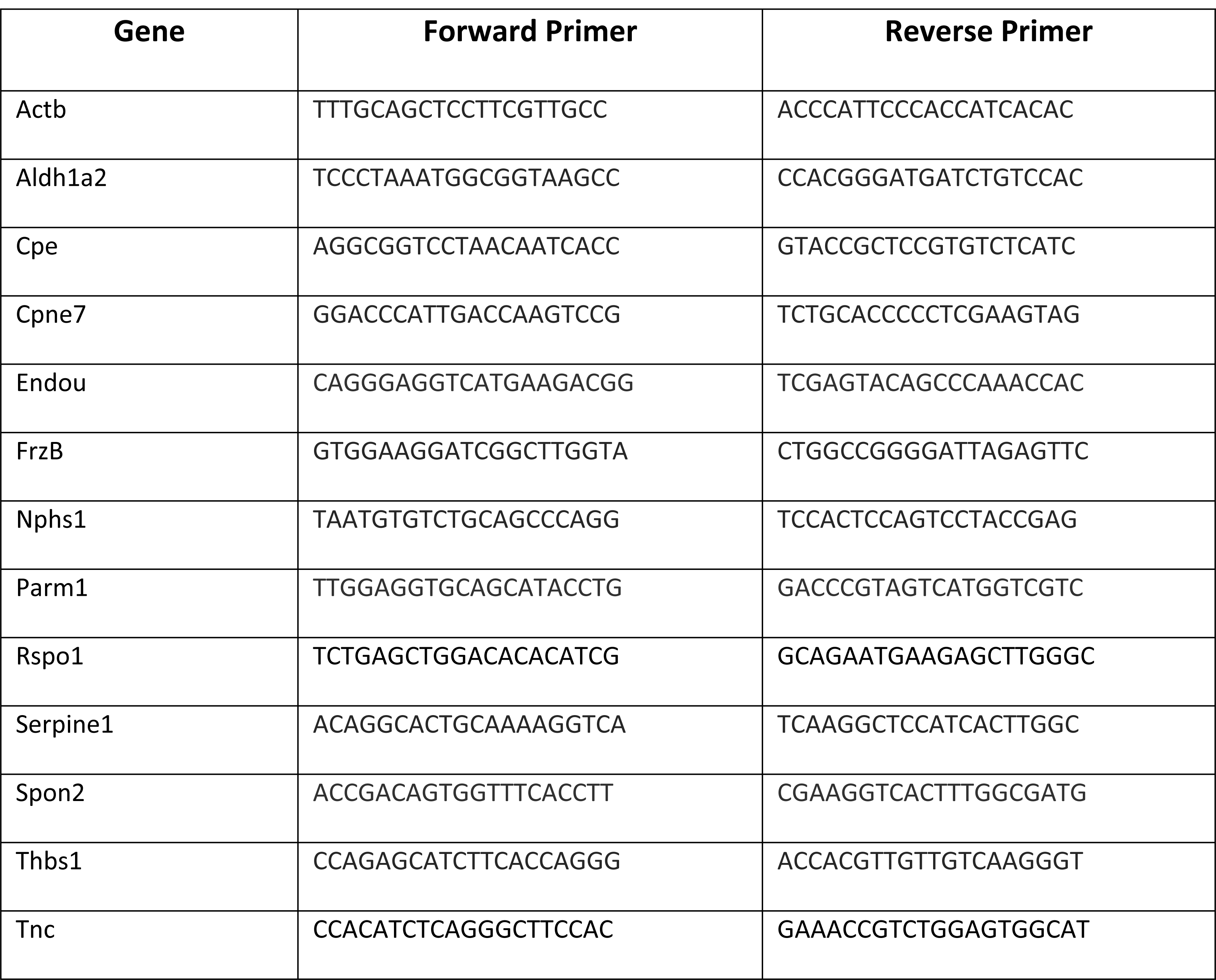
Primer sequences used for RT-qPCR validations.

### RNA-seq Data analysis

RNA-seq data was analyzed using Strand NGS 3.2. The reads were aligned to mm9. Reads were quantified using DeSeq with normalization threshold of 1. The baseline was set to the median of all samples.

### Filtering gene lists (all samples)

Reads were filtered based on genic region; spliced, partial genic, exonic, exon intron junction and genic reads were retained; intergenic and intronic reads were removed. Reads were further filtered on read quality metrics with the following parameters: Quality threshold >= 20, N’s allowed in read <=0, number of multiple matches allowed <= 1, and reads were removed that are failing vendor’s QC. Duplicate reads were removed, with a threshold of 4.

### Generating gene lists comparing podocytes, mesangial cells, endothelial cells and controls

At least 5.0 NRPKM in 3 out of 6 samples was required. A moderated t-test was then used to compare 3-allele and control samples with a corrected p-value cut-off of 0.05. The p-value computation was asymptotic and no multiple testing correction was used. This list was then used to perform a fold-change analysis between 3-allele and control samples, with a fold-change cut-off of >= 1.5. Y-linked genes, associated with sex differences between the samples, were removed from the final gene list.

### Filtering endothelial genes from MafB-GFP podocyte samples

For the podocytes, in order to remove the effects of a small percentage of contaminating endothelial cells, the WT glomerular endothelial cells were compared with WT podocytes. At least 80 NRPKM was required in 2 out of 6 samples (podocyte or endothelial); the cell types were compared with Audic Claverie test with no multiple-testing correction and a p-value cut off of 0.05. A fold change>=4 in endothelial cells compared to podocytes was used to generate a list of genes with high specific expression in endothelial cells. This list of genes was compared using a Venn diagram with differentially expressed podocyte (MAFB) genes. Genes were excluded from the list of differentially expressed MAFB genes which were found in both the MAFB list and the list of genes up-regulated in endothelial cells.

### Heat map parameters

The heat map was generated within Strand NGS software using a hierarchical clustering algorithm and clustered by normalized intensity values. The heat map was clustered on entities and conditions. The similarity measure is Pearson Centered. The linkage rule is Ward’s. There was no clustering within conditions.

## Results and Discussion

In this study we used RNA-seq to define the altered gene expression patterns of each major cell type of the glomerulus in the *Cd2ap*+/-, *Fyn*-/- bigenic mouse model of FSGS. To isolate the podocytes, mesangial cells and endothelial cells from the control and mutant FSGS kidneys we used three different transgenic reporter mouse lines. *MafB-GFP, Meis1-GFP* and *Tie2-GFP* which drive cell type restricted GFP expression in the podocytes, mesangial cells and endothelial cells of the glomerulus, respectively (24). We first isolated glomeruli, using a sieving protocol, and then used further enzymatic dissociation to produce single cell suspensions. By combining the three mutant alleles (*Cd2ap*+/-, *Fyn*-/-) with the appropriate GFP transgene reporter it was possible to purify the podocytes, mesangial cells and endothelial cells by fluorescent activated cell sorting (FACS). We then examined the FSGS altered gene expression patterns of all three major glomerular cell types using RNA-seq to begin to better understand the underlying pathogenic molecular pathways.

### *Cd2ap* +/-, *Fyn*-/- Phenotype

In agreement with previous studies, we observed the onset of proteinuria at around 5 months of age in the *Cd2ap*+/-, *Fyn*-/- mice, with near 100% penetrance (17). H&E and Jones Silver Stain were used to characterize the FSGS like pathology observed in the *Cd2ap*+/-, *Fyn*-/- glomeruli (Fig. 1). There were localized regions of scarring in some glomeruli, as well as capillary lumen loss and increased mesangial matrix. Scanning EM of the *Cd2ap*+/-, *Fyn*-/- glomeruli showed podocyte depletion, fusion of foot processes and general disorganization of remaining podocytes (Fig. 2). Transmission EM further showed the altered podocyte foot process structures and, of interest, regions of expanded GBM (Fig. 3). At the time of sacrifice, around 11 months, the *Cd2ap*+/-, *Fyn*-/- mice showed normal serum creatinine, elevated serum BUN (33.6 mg/dL versus 26.2 mg/dL for control, P = 0.0001), and decreased serum albumin (2.58 g/dL versus 2.89 g/dL for control, P = 0.03).

**Fig. 1.**
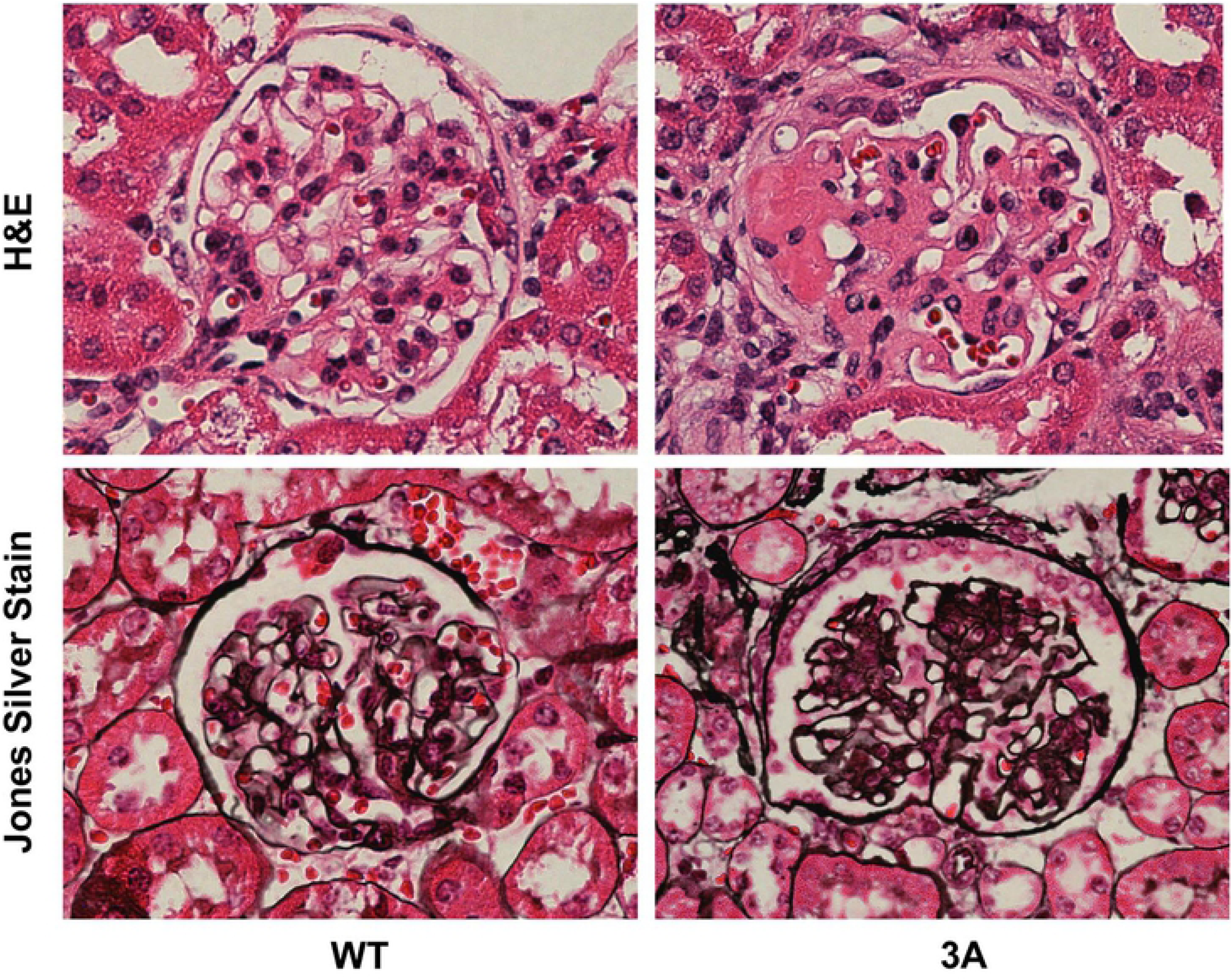
Histology of wild type (WT) and *Cd2ap*+/-, *Fyn*-/- three allele mutant (3A) glomeruli. H&E staining, top panels, show partial blockage of mutant capillaries and regions of scarring, particularly pronounced on the left extreme. Bottom panels show Jone’s Silver stain, with increased staining of glomerular basement membranes in 3A glomerulus.

**Fig. 2.**
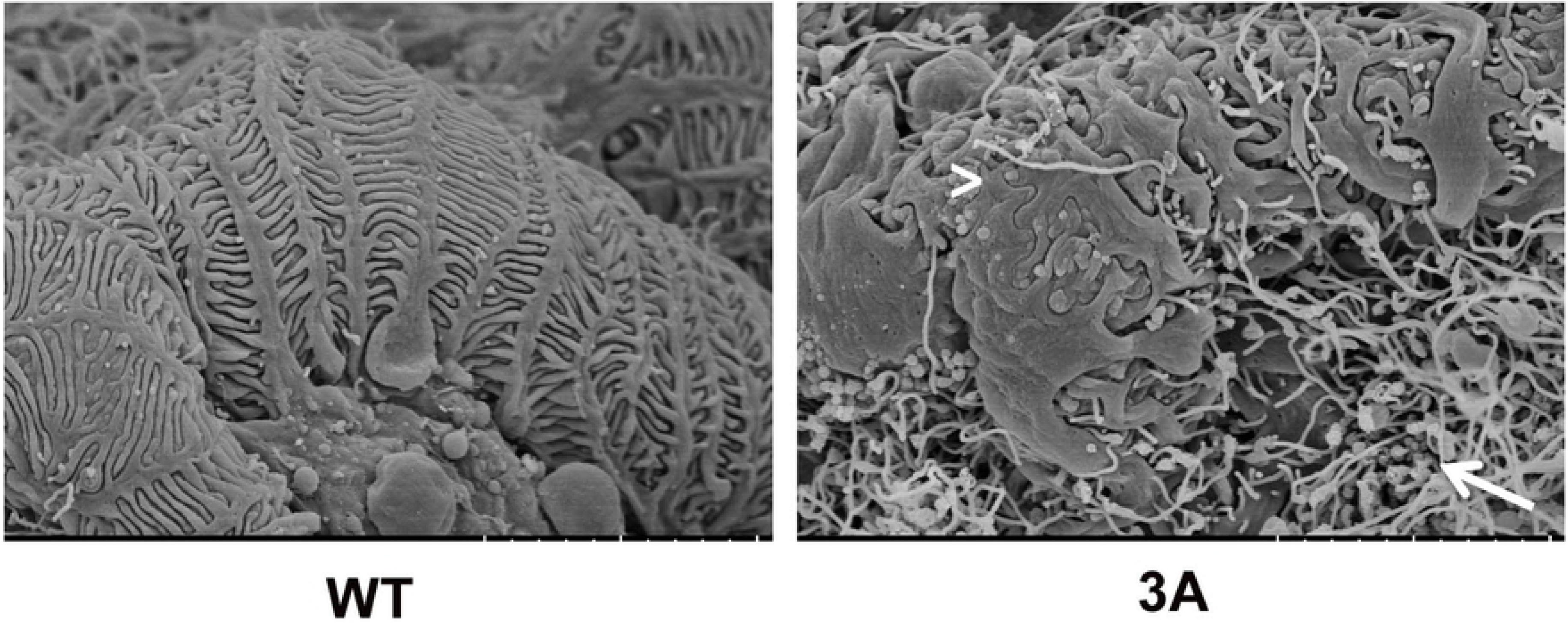
Scanning electron microscopy of wild type (WT) and three allele mutant (3A) glomeruli. Note uniform capillary coverage by wild type podocytes, with evenly spaced foot processes and slit diaphragms. In contrast the mutant podocytes are disorganized, partially detached, with foot processes sometimes fused (arrowhead), blocking slit diaphragms. In addition, the mutant glomerulus showed abundant fibrils (arrow).

**Fig. 3.**
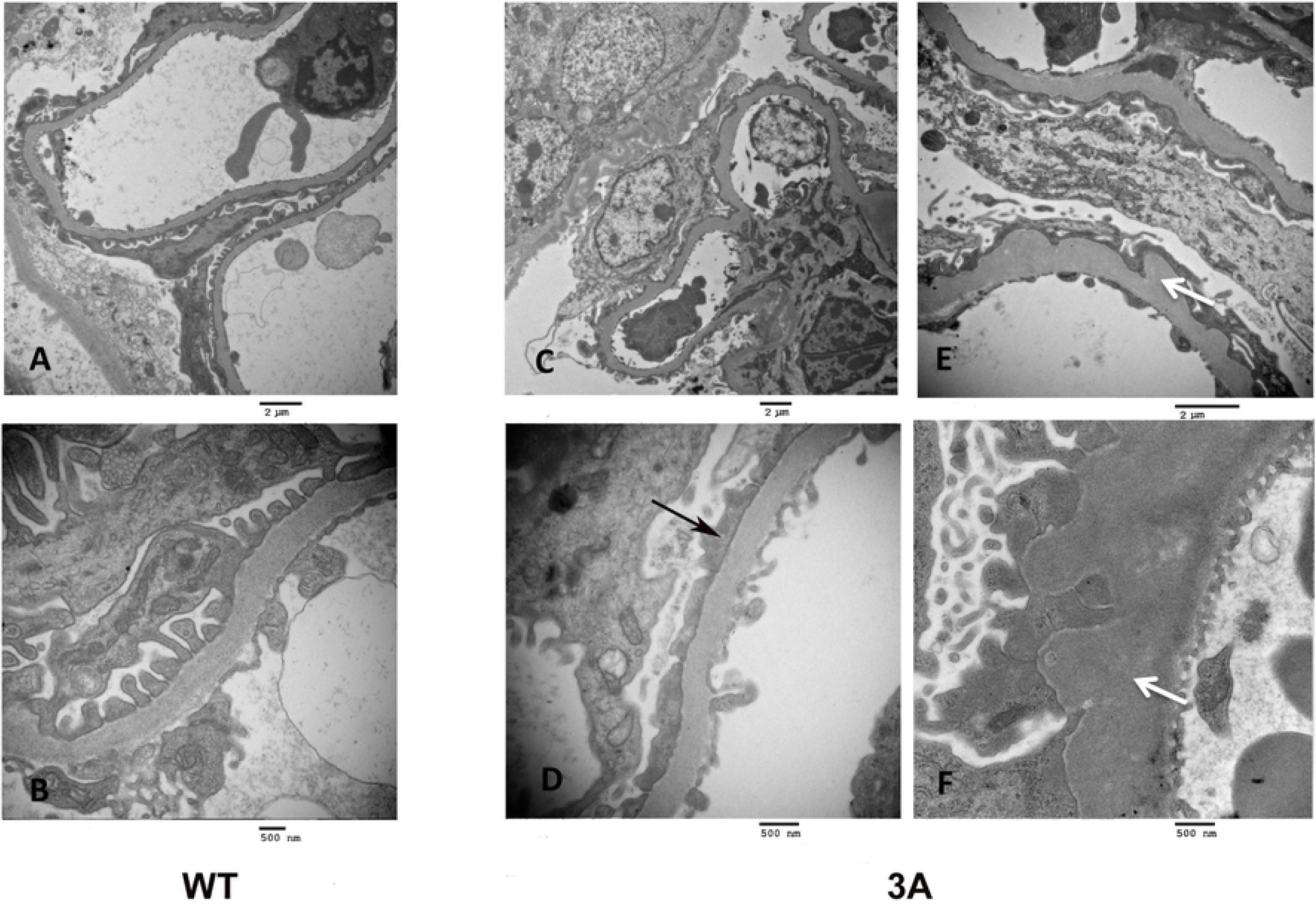
Transmission electron microscopy of glomeruli of wild type (WT), left panels, and mutant (3A), middle and right panels. Upper panels are lower magnification and lower panels are higher magnification (see size bars). Note evenly spaced foot processes and slit diaphragms in the WT, with the mutant showing fused foot processes (black arrow) and expanded glomerular basement membrane (white arrow).

### Assaying cell type purity

The isolation of strongly enriched populations of glomerular cell types can be challenging. The podocytes, in particular, are tightly wrapped around capillary loops with extensively interdigitated foot processes attached to the glomerular basement membrane. The mutant podocytes, within partially sclerotic glomeruli, were particularly challenging to detach.

We used two metrics to assay enrichment levels for the three cell types following FACS. The first was enriched expression levels of expected cell type specific marker genes. By this measure all FACS cell preparations showed strongly elevated expression of predicted markers, indicating robust enrichment of desired cell types. For example, *Nphs2* expression is an excellent marker for the podocyte, and all six podocyte cell preparations, including three *Cd2ap*+/-, *Fyn*-/- mutant and three control, showed very high expression levels of *Nphs2*, in the range of 1,400-2,300 RPKM.

A second and more stringent metric is to look for the expression levels of marker genes associated with potential contaminating cell types. By this measure, for example, the endothelial and mesangial cell preparations were very free of podocyte contamination, with podocyte marker *Nphs2* expression levels of only 0, 0, 0, 0, 4 and 1 RPKM in the six endothelial cell preparations and 0, 2, 0, 1, 1, and 11 RPKM in the mesangial cell preparations. In similar manner the mesangial cells were essentially free of endothelial contamination and the endothelial cells also free of mesangial contamination.

The podocytes, however, showed detectable levels of cross contamination. For example, the endothelial expressed gene *Kdr*, gave an average expression level of 336 RPKM in endothelial cell preparations and 15 RPKM in podocytes, suggesting a low level of contamination. The bioinformatics analysis pipeline was therefore modified for podocytes to take this into account, as described in Material and Methods.

### Podocytes

As previously mentioned, to define the gene expression pattern of mutant podocytes we made *Cd2ap*+/-, *Fyn*-/- mice that also carried the *MafB-GFP* transgene, which gives GFP expression specifically in podocytes in the developing kidney (26). Glomeruli were purified, from mutants and controls, dissociated to give single cell suspensions, and podocytes isolated by FACS. RNA was purified and used for RNA-seq, and the resulting data analyzed with Strand NGS software.

There were many significant gene expression changes in the *Cd2ap*+/-, *Fyn*-/- podocytes, compared to controls, with 90 genes up-regulated greater than 2 fold change (FC) and 29 genes down-regulated (> 2 FC) (Table S1). A heatmap showing gene expression changes in bigenic mutant podocytes compared to controls is shown in Fig. 4, with an expandable version including gene names provided (Fig. S1).

**Fig. 4.**
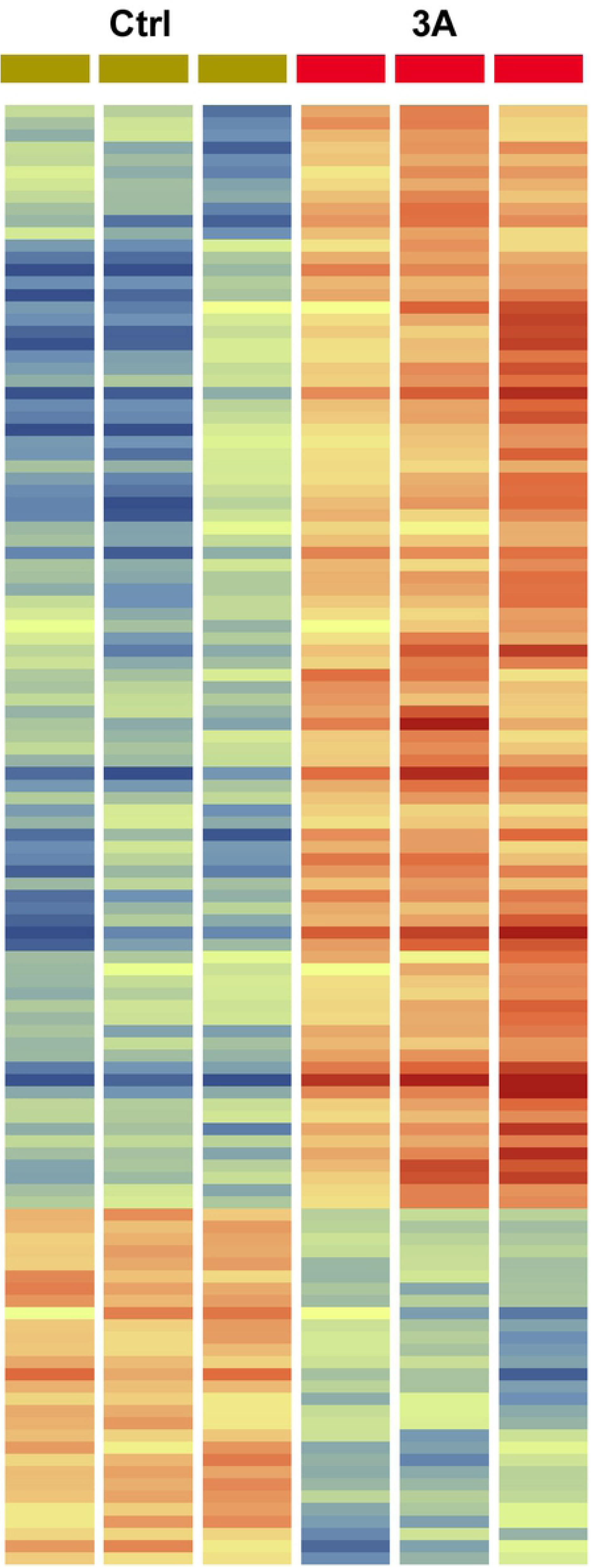
Heatmap showing gene expression changes in control (ctrl) versus *Cd2ap*+/-, *Fyn*-/- three allele mutant (3A) podocytes. There were three samples of each, with red indicating stronger expression and blue indicating weaker expression. Genes with greater than 2 fold change are shown. An expandable version including gene names is included in supplementary data (Fig. S1). Lists of genes with fold change are included in Table S1.

Growth factor related genes with elevated expression in mutant podocytes included *Gdnf* (22 FC), *Tnc* (22 FC), *Bmp4* (5 FC), *Tgfβ1* (2.5 FC), *Tgfβ2* (3.5 FC), *Cxcl12* (1.7 FC), *Nrp2* (3.0 FC) *Rspo1* (3.0 FC), *Ctgf* (1.5 FC), and *Vegfb* (2.0 FC). This group of genes includes a very strong Tgfβ family signature, as detailed below.

*Gdnf*, glial derived neurotrophic factor, showed 22 FC increased expression in podocytes of mutants. During kidney development Gdnf is expressed by cap mesenchyme nephron progenitors as well as stromal cells (27), driving branching morphogenesis of the ureteric bud, which expresses the Gdnf receptor Ret. Gdnf is a member of the Tgfβ superfamily. Of particular interest, Gdnf has been shown to be a survival factor in injured podocytes, acting in an autocrine manner (28).

The increased expression of *Tgfβ1, Tgfβ2, Vegfb* and *Bmp4*, additional members of the Tgfβ family, give evidence for a role for podocytes in driving fibrosis, since Tgfβ has been shown to be a major mediator of fibrosis (29, 30). *Bmp4* has been shown to play a profibrotic role in the liver (31), and to drive expression of the fibrosis related gene *Acta2* in hepatocyte stellate cells (32). Of note, we did also observe in the mutant podocytes strong upregulation (8.2 FC) of *Acta2*, a classic smooth muscle marker associated with myofibroblasts. During fibrosis many cell types can differentiate into myofibroblasts (33).

We also observed the upregulation of *Arid5b* (1.6 FC) in mutant podocytes. This gene encodes a component of a histone demethylase complex, resulting in target gene activation. Overexpression of *Arid5b* in fibroblasts can result in induction of smooth muscle genes, including smooth muscle actin, *Acta2* (34).

Also connected to the Tgfβ pathway, we observed upregulation of *Snai2* (5.8 FC), encoding the transcriptional repressor Snail. Tgfβ has been shown to induce Snail expression (35). Further, expression of Snail has been shown to activate Tgfβ signaling in breast cancer cells (36), providing another feed forward loop for this pathogenic pathway.

Tenascin-C (*Tnc*) showed ∼22 fold elevated expression in mutant podocytes. Tenascin-C is a multifunctional extracellular matrix protein that is upregulated during wound healing, with persistent expression observed in a variety of chronic pathological conditions (37). Importantly, *Tnc* expression in fibroblasts drives a fibrotic response, once again including upregulation of *Acta2* expression (38). Further, there is a dramatic attenuation of lung fibrosis in bleomycin treated mice carrying homozygous mutation of *Tnc* (38). *Tnc* expression is upregulated by Tgfβ in fibroblasts through Alk5 mediated Smad2/3 signaling (38). Tnc promotes secretion of Tgfβ, which in turn promotes expression of Tnc, resulting in yet another a feed forward loop (38).

*Tnc* transcripts undergo extensive alternative splicing, potentially giving rise to hundreds of different isoforms (39). In addition, the Tnc protein undergoes many variable post translational modifications, further contributing to its observed considerable functional diversity. Tnc can bind multiple ligands including fibronectin, contactin, glypican, and integrins (39), and can also activate epidermal grown factor receptor (40) and the Toll like receptor TLR-4 (41). Tnc is a poor adhesive substrate and blocks fibronectin mediated adhesion and is therefore anti-adhesive. The dramatic upregulation of *Tnc* in the mutant podocytes and its multiple functions including pro-fibrotic and anti-adhesive suggest a significant role in the disease process.

The observed upregulation of *Rspo1* (3 FC) is also of interest in relation to fibrosis/sclerosis. R-spondins are extracellular Wnt agonists that promote Wnt signaling by reducing degradation of Frizzled and LRP co-receptors (42). Further, canonical Wnt signaling has been shown to play a central role in multiple fibrosing diseases, including pulmonary, renal and liver fibrosis (43-45).

The mutant podocytes also showed strong evidence of elevated retinoic acid (RA) related pathways. Expression of *Aldh1a1*, encoding an enzyme driving the final step of RA synthesis (46), was increased 7 fold in *Cd2ap*+/-, *Fyn*-/- mutants. RA can induce expression of *Ripply3* as well as *Tbx1* in the pre-placodal ectoderm (47). Retinoic acid signaling can also induce *Tbx3* during limb development (48). Of interest, we observed upregulation of both *Ripply3* (7 FC) and *Tbx3* (5.2 FC), in mutant podocytes, consistent with elevated RA signaling. Ripply3 is a transcriptional repressor that can interact with the transcription factor Tbx1, converting it from an activator to a repressor (47). In the adriamycin murine model of FSGS reducing RA synthesis resulted in increased proteinuria and glomerulosclerosis, while treatment with RA reduced proteinuria and reduced podocyte loss (49). The observed elevated RA synthesis in the *Cd2ap*+/-, *Fyn*-/- mutant podocytes can therefore be viewed as a protective response.

Another highly upregulated gene in the mutant podocytes was *Spon2* (135 FC), which encodes an extracellular matrix protein that is critical for inflammatory cell recruitment (50), efficient dendritic cell priming of T-lymphocytes (51) and trafficking of eosinophils (52), suggesting an important role in driving the immune response. *Fxyd5* (9.6 FC), encoding the gamma subunit of the Na,K-ATPase, as well as *Atp1b1* (9.4 FC), encoding the beta chain of Na,K-ATPase were also increased in expression in mutant podocytes. *Sost*, encoding sclerostin, was also strongly upregulated (10 FC) in mutant podocytes. It is a BMP antagonist, perhaps counteracting in some measure the observed upregulation of Bmp4.

GO analysis of the list of genes upregulated in the mutant podocytes identified elevated protein kinase activity and increased expression of growth factors as the most significant molecular functions. The most strongly impacted biological process was negative regulation of cell adhesion (P = 4.26E-9), likely related to FSGS loss of podocytes. Other upregulated biological processes were cell motility (P = 7.8E-9) and programmed cell death (P = 3.5E-8). The observed apoptosis gene expression signature in the mutant podocytes, with for example elevated expression of *Casp4* (5.8 FC), is also consistent with the well-defined podocyte depletion in FSGS (53, 54). Fig. 5 shows a cytoscape with multiple identified functionalities and associated genes.

**Fig. 5.**
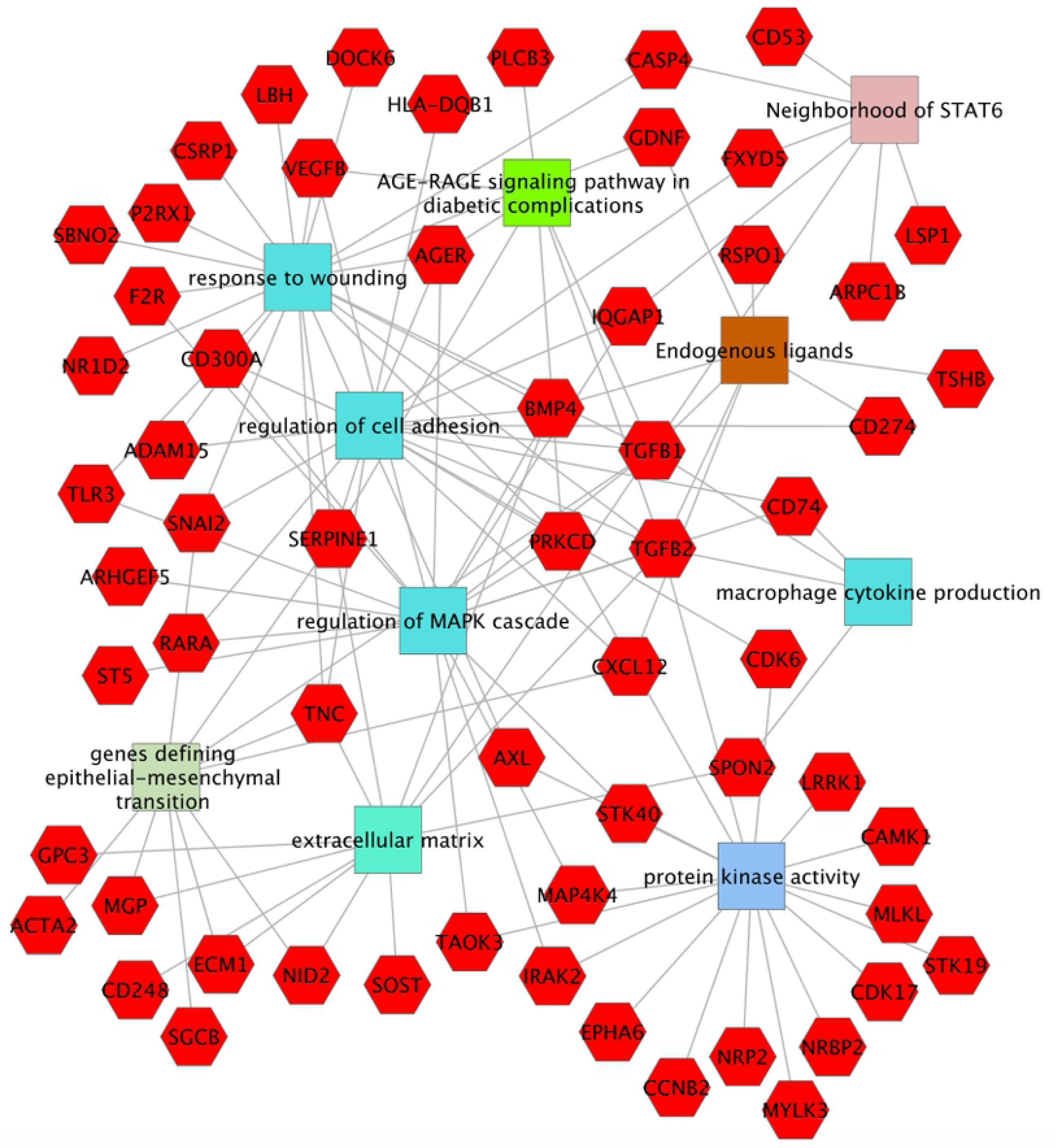
Cytoscape showing some of the genes upregulated in *Cd2ap*+/-, *Fyn*-/- mutant podocytes and their associated functionalities. There were fewer genes showing strong down regulation compared to upregulation for the mutant podocytes. Among downregulated genes were *Robo2* (1.5 FC) and *Dpysl2* (1.6 FC), both involved in axon guidance, and *Ncam1* (3.2 FC), encoding neural cell adhesion molecule, all perhaps reflecting a reduced neural character of the mutant podocytes. Also of interest, GO analysis of the list of down regulated genes gave the strongest signature for reduced protein ubiquitination (uncorrected P=3.4E-5), with seventeen associated genes.

The *Actn4*-/- genetic murine model of FSGS was previously transcriptionally profiled using the Translating Ribosome Affinity Purification (TRAP) method, with cell type specific expression of transgenic Collagen-1α1-eGFP-L10a allowing affinity enrichment of podocyte expressed RNAs (55). Of interest, the podocytes of *Actn4*-/- mice also showed elevated expression of *aldehyde dehydrogenase*, similar to the *Cd2ap*+/-, *Fyn*-/- mutant podocytes in this report. Further, the podocytes of *Actn4-/-* mice showed increased expression of *Pamr1*, which also gave elevated expression in the *Cd2ap*+/-, *Fyn*-/- mutant podocytes (3.6 FC, Table S1). The *Pamr1* gene has been associated with muscle regeneration (56), which is of interest given the contractile nature of the podocyte (57), perhaps playing a role in counteracting the perfusion pressure of the capillaries. Nevertheless, the gene expression signature of the podocyte shows only a weak muscle signature (26).

### Mesangial cells

Mesangial expansion is a hallmark of FSGS, with increased extracellular matrix and mesangial cell proliferation. The *Cd2ap*+/-, *Fyn*-/- mutant mesangial cells showed upregulation of 55 genes (>1.5 FC) compared to control (Table S2). One of the upregulated genes (4.3 FC) with strong expression, over 100 RPKM in mutants, was *Aldh1a2*, involved in the synthesis of retinoic acid, similar to *Aldh1a1*, which was upregulated in podocytes.

Thrombospondin (*Thbs1*, 3.8 FC) was strongly upregulated in mutant mesangial cells. Thbs plays an important role in the activation of Tgfβ, which is secreted in inactive pro-cytokine form. The inflammatory phenotype of *Thbs1* mutants closely resembles that of *Tgfβ* mutants (58). Given the key role of Tgfβ in fibrosis the elevated *Thbs1* expression in mutant mesangial cells is likely pro-fibrotic.

Upregulated mesangial cell genes in *Cd2ap*+/-, *Fyn*-/- mutants also included *ccdc68* (3.9 FC), encoding a centriole protein, *Frzb* (4.1 FC) encoding a secreted Wnt binding protein involved in the regulation of bone development, *Tnnt2* (4.1 FC) involved in muscle contraction, *Col8a1* (1.9 FC) involved in extracellular matrix, and *F2r* (1.6 FC), encoding a G-protein coupled thrombin receptor involved in the thrombotic response.

It is interesting to again compare the altered gene expression observed in *Cd2ap*+/-, *Fyn*-/- mutant mesangial cells in this report with previously observed gene expression changes in other mouse genetic models of kidney disease. Mesangial cell upregulated genes in *Cd2ap*+/-, *Fyn*-/- mice that were also upregulated in the mesangial cells of the *db/db* diabetic nephropathy mouse model include *Cpe, Anxa3, Thbs1, Pmp22, Tspan2* and *Akap12*, and closely related gene family members for *Ppap2c, Mmp3* and *Adam22* (59). Some of the top genes upregulated in *Cd2ap*-/- mesangial cells, and also upregulated in *Cd2ap*+/-, *Fyn*-/- mutant mesangial cells include *Col8a1, Ccdc68, Thbs1, Tnnt2, Frzb, Pmp22*, and *Aldh1a2* (24). The *Thbs1* gene stands out as strongly upregulated in all three genetic models. The robust correlation for *Cd2ap*-/- and *Cd2ap*+/-, *Fyn*-/- mutant mesangial cells is perhaps not surprising given the overlapping genetics.

It was interesting to find that in some cases the same pathway was activated in both podocytes and mesangial cells. For example, aldehyde dehydrogenase activity, reflecting increased RA synthesis, was elevated in both *Cd2ap*+/-, *Fyn*-/- podocytes and mesangial cells. Further, *Prss23* (4.7 FC) was one of the most strongly upregulated genes in *Cd2ap*+/-, *Fyn*-/- mesangial cells, and also one of the most strongly upregulated genes in podocytes of the *Cd2ap*-/- FSGS model (24), although, interestingly, it was not strongly upregulated in *Cd2ap*+/-, *Fyn*-/- podocytes. In addition, *Prss23* is upregulated in the glomeruli of human patients with FSGS (60). This common thread of upregulated *Prss23* in multiple mouse models, and cell types, as well as in human FSGS gives evidence for a significant role. The serine protease encoded by *Prss23* can activate Par2 (Protease-Activated Receptor 2), which has been implicated in TGFβ1 induced podocyte injury in the Adriamycin model of nephropathy in rats (61). It has also been shown that Prss23 can promote TGFβ signaling during zebrafish cardiac valve formation as well as in human aortic endothelial cell assays (62).

There were relatively few down regulated genes in *Cd2ap*+/-, *Fyn*-/- mutant mesangial cells, with only six with greater than 2 FC, and all of these showing low expression levels, even in controls (Table S2).

### Endothelial cells

The endothelial cells are the third major cell type of the glomerulus. They can also play a role in the progression of FSGS through the production of growth factors and cytokines, leaky angiogenesis, and the recruitment of macrophages and leukocytes. The observed gene expression changes in the *Cd2ap*+/-, *Fyn*-/- mutant endothelial cells were modest in number, with only 45 genes showing greater than 1.5 and 15 genes with greater than 2 FC upregulation. There were even fewer downregulated genes in mutants, with 26 showing >1.5 FC and only 5 with >2.0 FC (Table S3).

Upregulated genes included *Apnlr* encoding a G-coupled receptor for Apelin that inhibits adenylate cyclase and is implicated in angioblast migration and regulation of blood vessel formation. Also upregulated was *Kctd10*, which binds proliferating cell nuclear antigen (PCNA) and may be involved in DNA synthesis and cell proliferation and is also involved in ubiquitination.

Another upregulated gene in mutant endothelial cells was *Nestin*, which encodes an intermediate filament protein and is normally associated with neural stem/progenitor cells. It is, however, also expressed in endothelial cells in pancreas (63), in ovary and placenta (64), as well as vascular neoplasms (65) and cancers (66), where is a marker of neovasculature.

GO analysis for mutant endothelial cell upregulated genes found biological processes including negative regulation of response to oxidative stress, negative regulation of cell death and lymphocyte aggregation.

### QPCR Validations

The RNA-seq results were validated using RT-QPCR. Glomeruli from three mutant allele *Cd2ap*+/-, *Fyn*-/- and control mice were isolated using a sieving procedure (24), RNA purified and used for RT-QPCR. Genes were selected based on restricted up-regulation in mutants in one sorted glomerular cell type compared to control cells. Samples were normalized to beta actin (*actb*) for mesangial and endothelial genes, or nephrin (*Nphs2*) for podocyte genes. RT-QPCR validated upregulation of *Endou, Parm1, Serpine1, Spon2*, and *Rspo1* in mutant podocytes, and *Cpe, Aldh1a2, Tnc, Cpne7, Frzb*, and *Thbs1* in mutant mesangial cells (Fig. 6).

**Fig. 6.**
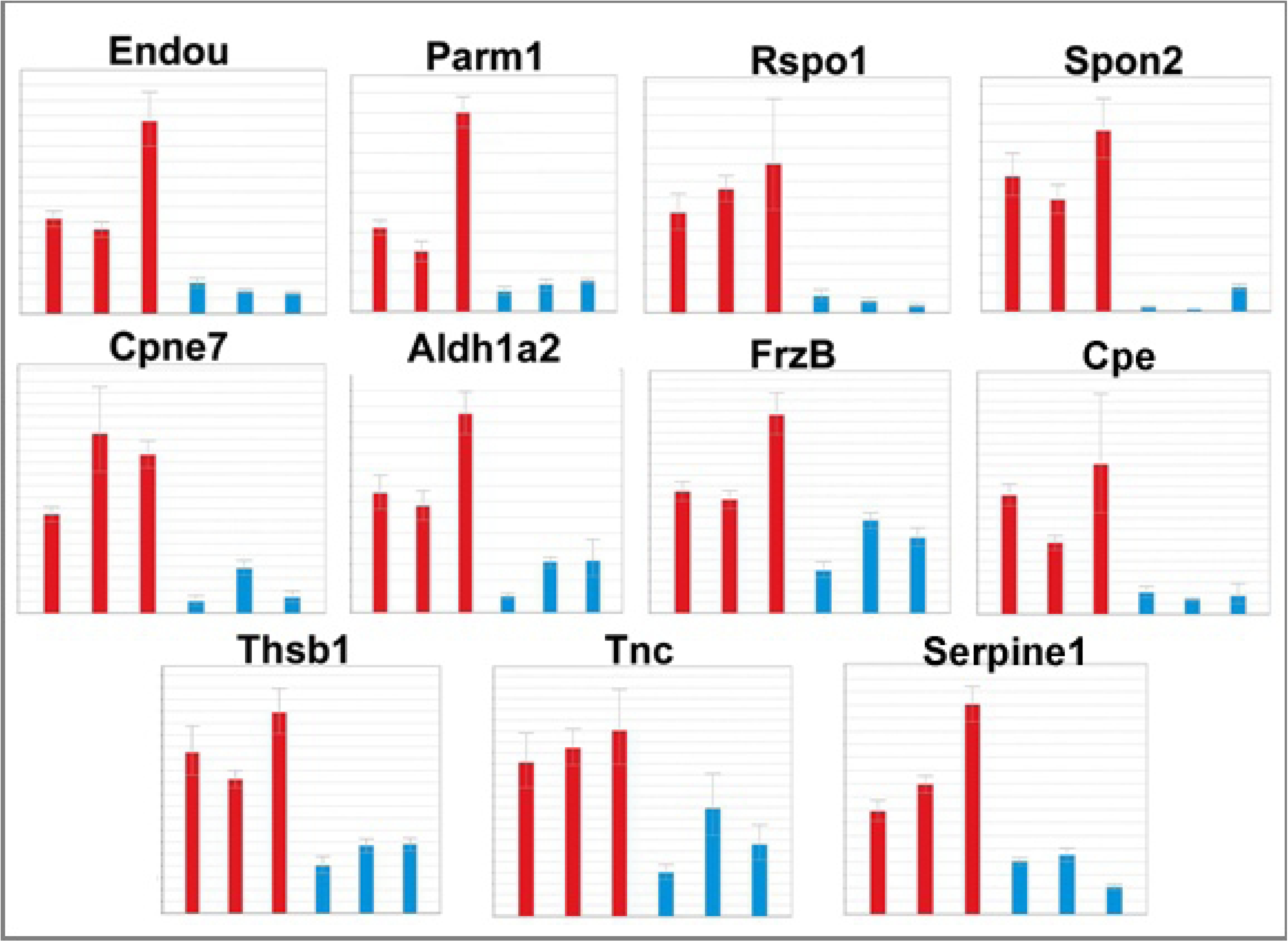

## Conclusions

FSGS is a major cause of ESRD, with a high percentage of patients unresponsive to available therapies. Improved understanding of the molecular underpinnings of the disease process could provide insights leading to novel therapeutic approaches. In this study we carry out an RNA-seq analysis of the altered gene expression patterns of podocytes, mesangial cells and glomerular endothelial cells of the bigenic *Cd2ap*+/-,

*Fyn*-/- mutant mouse model of FSGS. The podocytes showed the most dramatic changes, with upregulation of many genes related to the Tgfβ family/pathway, including *Gdnf, Tgfβ1, Tgfβ2, Snai2, Vegfb, Bmp4*, and *Tnc*. The mutant podocytes also showed upregulation of *Acta2*, a marker of smooth muscle and associated with myofibroblasts, which are implicated in driving fibrosis. GO analysis of the podocyte upregulated genes identified elevated protein kinase activity, increased expression of growth factors, and negative regulation of cell adhesion, perhaps related to the observed podocyte loss in FSGS.

Both podocytes and mesangial cells showed strong upregulation of aldehyde dehydrogenase genes involved in the synthesis of RA. Similarly, the *Cd2ap*+/-, *Fyn*-/- mesangial cells, as well as podocytes in other genetic models, and the glomeruli of human FSGS patients, all show upregulation of the serine protease Prss23, with the common thread suggesting important functionality. Another gene with strong upregulation in the *Cd2ap*+/-, *Fyn*-/- mutant mesangial cells as well as multiple other mutant mouse models of FSGS was thrombospondin, which activates the secreted inactive form of Tgfβ.

The *Cd2ap*+/-, *Fyn*-/- mutant endothelial cells showed elevated expression of genes involved in cell proliferation, angioblast migration, angiogenesis, and neovasculature, all consistent with the formation of new blood vessels in the diseased glomerulus. In total the data herein provide a global definition of the pathogenic and protective molecular pathways that are activated in the three major cell types of the glomerulus in the bigenic *Cd2ap*+/-, *Fyn*-/- mouse model of FSGS.

## Acknowledgments

We thank Hung Chi Liang for carrying out amplification reactions for RNA-seq analysis. We thank Dominic Distasio for assistance in managing the mouse colony. This work was supported by the National Institutes of Health grant P50 DK096418.

## References

1. Kitiyakara C, Kopp JB, Eggers P. Trends in the epidemiology of focal segmental glomerulosclerosis. Semin Nephrol. 2003;23(2):172–82.

2. Sprangers B, Meijers B, Appel G. FSGS: Diagnosis and Diagnostic Work-Up. Biomed Res Int. 2016;2016:4632768.

3. Kitiyakara C, Eggers P, Kopp JB. Twenty-one-year trend in ESRD due to focal segmental glomerulosclerosis in the United States. Am J Kidney Dis. 2004;44(5):815–25.

4. Asinobi AO, Ademola AD, Okolo CA, Yaria JO. Trends in the histopathology of childhood nephrotic syndrome in Ibadan Nigeria: preponderance of idiopathic focal segmental glomerulosclerosis. BMC Nephrol. 2015;16:213.

5. D’Agati VD, Fogo AB, Bruijn JA, Jennette JC. Pathologic classification of focal segmental glomerulosclerosis: a working proposal. Am J Kidney Dis. 2004;43(2):368–82.

6. Meehan SM, Chang A, Gibson IW, Kim L, Kambham N, Laszik Z. A study of interobserver reproducibility of morphologic lesions of focal segmental glomerulosclerosis. Virchows Arch. 2013;462(2):229–37.

7. Rosenberg AZ, Kopp JB. Focal Segmental Glomerulosclerosis. Clin J Am Soc Nephrol. 2017;12(3):502–17.

8. Sadowski CE, Lovric S, Ashraf S, Pabst WL, Gee HY, Kohl S, et al. A single-gene cause in 29.5% of cases of steroid-resistant nephrotic syndrome. J Am Soc Nephrol. 2015;26(6):1279–89.

9. Trautmann A, Bodria M, Ozaltin F, Gheisari A, Melk A, Azocar M, et al. Spectrum of steroid-resistant and congenital nephrotic syndrome in children: the PodoNet registry cohort. Clin J Am Soc Nephrol. 2015;10(4):592–600.

10. Kopp JB, Nelson GW, Sampath K, Johnson RC, Genovese G, An P, et al. APOL1 genetic variants in focal segmental glomerulosclerosis and HIV-associated nephropathy. J Am Soc Nephrol. 2011;22(11):2129–37.

11. Mele C, Iatropoulos P, Donadelli R, Calabria A, Maranta R, Cassis P, et al. MYO1E mutations and childhood familial focal segmental glomerulosclerosis. N Engl J Med. 2011;365(4):295–306.

12. Lennon R, Stuart HM, Bierzynska A, Randles MJ, Kerr B, Hillman KA, et al. Coinheritance of COL4A5 and MYO1E mutations accentuate the severity of kidney disease. Pediatr Nephrol. 2015;30(9):1459–65.

13. Shih NY, Li J, Cotran R, Mundel P, Miner JH, Shaw AS. CD2AP localizes to the slit diaphragm and binds to nephrin via a novel C-terminal domain. Am J Pathol. 2001;159(6):2303–8.

14. Schwarz K, Simons M, Reiser J, Saleem MA, Faul C, Kriz W, et al. Podocin, a raft-associated component of the glomerular slit diaphragm, interacts with CD2AP and nephrin. J Clin Invest. 2001;108(11):1621–9.

15. Gigante M, Pontrelli P, Montemurno E, Roca L, Aucella F, Penza R, et al. CD2AP mutations are associated with sporadic nephrotic syndrome and focal segmental glomerulosclerosis (FSGS). Nephrol Dial Transplant. 2009;24(6):1858–64.

16. Lowik MM, Groenen PJ, Pronk I, Lilien MR, Goldschmeding R, Dijkman HB, et al. Focal segmental glomerulosclerosis in a patient homozygous for a CD2AP mutation. Kidney Int. 2007;72(10):1198–203.

17. Huber TB, Kwoh C, Wu H, Asanuma K, Godel M, Hartleben B, et al. Bigenic mouse models of focal segmental glomerulosclerosis involving pairwise interaction of CD2AP, Fyn, and synaptopodin. J Clin Invest. 2006;116(5):1337–45.

18. Kim JM, Wu H, Green G, Winkler CA, Kopp JB, Miner JH, et al. CD2-associated protein haploinsufficiency is linked to glomerular disease susceptibility. Science. 2003;300(5623):1298–300.

19. Verma R, Wharram B, Kovari I, Kunkel R, Nihalani D, Wary KK, et al. Fyn binds to and phosphorylates the kidney slit diaphragm component Nephrin. J Biol Chem. 2003;278(23):20716–23.

20. Reiser J, Altintas MM. Podocytes. F1000Res. 2016;5.

21. Lim BJ, Yang JW, Do WS, Fogo AB. Pathogenesis of Focal Segmental Glomerulosclerosis. J Pathol Transl Med. 2016;50(6):405–10.

22. Galkina E, Ley K. Leukocyte recruitment and vascular injury in diabetic nephropathy. J Am Soc Nephrol. 2006;17(2):368–77.

23. Zent R, Pozzi A. Angiogenesis in diabetic nephropathy. Semin Nephrol. 2007;27(2):161–71.

24. Brunskill EW, Potter SS. Pathogenic pathways are activated in each major cell type of the glomerulus in the Cd2ap mutant mouse model of focal segmental glomerulosclerosis. BMC Nephrol. 2015;16:71.

25. Boerries M, Grahammer F, Eiselein S, Buck M, Meyer C, Goedel M, et al. Molecular fingerprinting of the podocyte reveals novel gene and protein regulatory networks. Kidney Int. 2013;83(6):1052–64.

26. Brunskill EW, Georgas K, Rumballe B, Little MH, Potter SS. Defining the molecular character of the developing and adult kidney podocyte. PLoS One. 2011;6(9):e24640.

27. Magella B, Adam M, Potter AS, Venkatasubramanian M, Chetal K, Hay SB, et al. Cross-platform single cell analysis of kidney development shows stromal cells express Gdnf. Dev Biol. 2018;434(1):36–47.

28. Tsui CC, Shankland SJ, Pierchala BA. Glial cell line-derived neurotrophic factor and its receptor ret is a novel ligand-receptor complex critical for survival response during podocyte injury. J Am Soc Nephrol. 2006;17(6):1543–52.

29. Branton MH, Kopp JB. TGF-beta and fibrosis. Microbes Infect. 1999;1(15):1349–65.

30. Biernacka A, Dobaczewski M, Frangogiannis NG. TGF-beta signaling in fibrosis. Growth Factors. 2011;29(5):196–202.

31. Fan J, Shen H, Sun Y, Li P, Burczynski F, Namaka M, et al. Bone morphogenetic protein 4 mediates bile duct ligation induced liver fibrosis through activation of Smad1 and ERK1/2 in rat hepatic stellate cells. J Cell Physiol. 2006;207(2):499–505.

32. Shen H, Huang G, Hadi M, Choy P, Zhang M, Minuk GY, et al. Transforming growth factor-beta1 downregulation of Smad1 gene expression in rat hepatic stellate cells. Am J Physiol Gastrointest Liver Physiol. 2003;285(3):G539–46.

33. Bergmann C, Distler JH. Canonical Wnt signaling in systemic sclerosis. Lab Invest. 2016;96(2):151–5.

34. Watanabe M, Layne MD, Hsieh CM, Maemura K, Gray S, Lee ME, et al. Regulation of smooth muscle cell differentiation by AT-rich interaction domain transcription factors Mrf2alpha and Mrf2beta. Circ Res. 2002;91(5):382–9.

35. Peinado H, Quintanilla M, Cano A. Transforming growth factor beta-1 induces snail transcription factor in epithelial cell lines: mechanisms for epithelial mesenchymal transitions. J Biol Chem. 2003;278(23):21113–23.

36. Dhasarathy A, Phadke D, Mav D, Shah RR, Wade PA. The transcription factors Snail and Slug activate the transforming growth factor-beta signaling pathway in breast cancer. PLoS One. 2011;6(10):e26514.

37. Udalova IA, Ruhmann M, Thomson SJ, Midwood KS. Expression and immune function of tenascin-C. Crit Rev Immunol. 2011;31(2):115–45.

38. Bhattacharyya S, Wang W, Morales-Nebreda L, Feng G, Wu M, Zhou X, et al. Tenascin-C drives persistence of organ fibrosis. Nat Commun. 2016;7:11703.

39. Midwood KS, Chiquet M, Tucker RP, Orend G. Tenascin-C at a glance. J Cell Sci. 2016;129(23):4321–7.

40. Swindle CS, Tran KT, Johnson TD, Banerjee P, Mayes AM, Griffith L, et al. Epidermal growth factor (EGF)-like repeats of human tenascin-C as ligands for EGF receptor. J Cell Biol. 2001;154(2):459–68.

41. Midwood KS, Orend G. The role of tenascin-C in tissue injury and tumorigenesis. J Cell Commun Signal. 2009;3(3-4):287-310.

42. Hao HX, Xie Y, Zhang Y, Charlat O, Oster E, Avello M, et al. ZNRF3 promotes Wnt receptor turnover in an R-spondin-sensitive manner. Nature. 2012;485(7397):195–200.

43. Akhmetshina A, Palumbo K, Dees C, Bergmann C, Venalis P, Zerr P, et al. Activation of canonical Wnt signalling is required for TGF-beta-mediated fibrosis. Nat Commun. 2012;3:735.

44. He W, Dai C, Li Y, Zeng G, Monga SP, Liu Y. Wnt/beta-catenin signaling promotes renal interstitial fibrosis. J Am Soc Nephrol. 2009;20(4):765–76.

45. Konigshoff M, Kramer M, Balsara N, Wilhelm J, Amarie OV, Jahn A, et al. WNT1-inducible signaling protein-1 mediates pulmonary fibrosis in mice and is upregulated in humans with idiopathic pulmonary fibrosis. J Clin Invest. 2009;119(4):772–87.

46. Bowles J, Feng CW, Miles K, Ineson J, Spiller C, Koopman P. ALDH1A1 provides a source of meiosis-inducing retinoic acid in mouse fetal ovaries. Nat Commun. 2016;7:10845.

47. Janesick A, Shiotsugu J, Taketani M, Blumberg B. RIPPLY3 is a retinoic acid-inducible repressor required for setting the borders of the pre-placodal ectoderm. Development. 2012;139(6):1213–24.

48. Ballim RD, Mendelsohn C, Papaioannou VE, Prince S. The ulnar-mammary syndrome gene, Tbx3, is a direct target of the retinoic acid signaling pathway, which regulates its expression during mouse limb development. Mol Biol Cell. 2012;23(12):2362–72.

49. Peired A, Angelotti ML, Ronconi E, la Marca G, Mazzinghi B, Sisti A, et al. Proteinuria impairs podocyte regeneration by sequestering retinoic acid. J Am Soc Nephrol. 2013;24(11):1756–68.

50. Jia W, Li H, He YW. The extracellular matrix protein mindin serves as an integrin ligand and is critical for inflammatory cell recruitment. Blood. 2005;106(12):3854–9.

51. Li H, Oliver T, Jia W, He YW. Efficient dendritic cell priming of T lymphocytes depends on the extracellular matrix protein mindin. EMBO J. 2006;25(17):4097–107.

52. Li Z, Garantziotis S, Jia W, Potts EN, Lalani S, Liu Z, et al. The extracellular matrix protein mindin regulates trafficking of murine eosinophils into the airspace. J Leukoc Biol. 2009;85(1):124–31.

53. Shankland SJ. The podocyte’s response to injury: role in proteinuria and glomerulosclerosis. Kidney Int. 2006;69(12):2131–47.

54. Jefferson JA, Shankland SJ. The pathogenesis of focal segmental glomerulosclerosis. Adv Chronic Kidney Dis. 2014;21(5):408–16.

55. Grgic I, Hofmeister AF, Genovese G, Bernhardy AJ, Sun H, Maarouf OH, et al. Discovery of new glomerular disease-relevant genes by translational profiling of podocytes in vivo. Kidney Int. 2014;86(6):1116–29.

56. Nakayama Y, Nara N, Kawakita Y, Takeshima Y, Arakawa M, Katoh M, et al. Cloning of cDNA encoding a regeneration-associated muscle protease whose expression is attenuated in cell lines derived from Duchenne muscular dystrophy patients. Am J Pathol. 2004;164(5):1773–82.

57. Andrews PM, Coffey AK. Cytoplasmic contractile elements in glomerular cells. Fed Proc. 1983;42(14):3046–52.

58. Crawford SE, Stellmach V, Murphy-Ullrich JE, Ribeiro SM, Lawler J, Hynes RO, et al. Thrombospondin-1 is a major activator of TGF-beta1 in vivo. Cell. 1998;93(7):1159–70.

59. Brunskill EW, Potter SS. Changes in the gene expression programs of renal mesangial cells during diabetic nephropathy. BMC Nephrol. 2012;13:70.

60. Bennett MR, Czech KA, Arend LJ, Witte DP, Devarajan P, Potter SS. Laser capture microdissection-microarray analysis of focal segmental glomerulosclerosis glomeruli. Nephron Exp Nephrol. 2007;107(1):e30–40.

61. Wang Y, He Y, Wang M, Lv P, Liu J, Wang J. Role of Protease-Activated Receptor 2 in Regulating Focal Segmental Glomerulosclerosis. Cell Physiol Biochem. 2017;41(3):1147–55.

62. Chen IH, Wang HH, Hsieh YS, Huang WC, Yeh HI, Chuang YJ. PRSS23 is essential for the Snail-ependent endothelial-to-mesenchymal transition during valvulogenesis in zebrafish. Cardiovasc Res. 2013;97(3):443–53.

63. Klein T, Ling Z, Heimberg H, Madsen OD, Heller RS, Serup P. Nestin is expressed in vascular endothelial cells in the adult human pancreas. J Histochem Cytochem. 2003;51(6):697–706.

64. Mokry J, Cizkova D, Filip S, Ehrmann J, Osterreicher J, Kolar Z, et al. Nestin expression by newly formed human blood vessels. Stem Cells Dev. 2004;13(6):658–64.

65. Shimizu T, Sugawara K, Tosaka M, Imai H, Hoya K, Takeuchi T, et al. Nestin expression in vascular malformations: a novel marker for proliferative endothelium. Neurol Med Chir (Tokyo). 2006;46(3):111–7.

66. Teranishi N, Naito Z, Ishiwata T, Tanaka N, Furukawa K, Seya T, et al. Identification of neovasculature using nestin in colorectal cancer. Int J Oncol. 2007;30(3):593–603.

